# Non-canonical function of the *Sex-lethal* gene controls the protogyny phenotype in *Drosophila melanogaster*

**DOI:** 10.1101/2021.03.02.433534

**Authors:** Ki-Hyeon Seong, Siu Kang

**Affiliations:** RIKEN Cluster for Pioneering Research, RIKEN Tsukuba Institute, Tsukuba, Ibaraki, Japan; Graduate School of Science and Engineering, Yamagata University, Jonan, Yonezawa, Yamagata, Japan; AMED-CREST, AMED, Chiyoda-ku, Tokyo, Japan

## Abstract

Many animal species exhibit sex differences in the time period prior to reaching sexual maturity. However, the underlying mechanism for such biased maturation remains poorly understood. Females of the fruit fly *Drosophila melanogaster* eclose 4 h faster on average than males, owing to differences in the pupal period between the sexes; this characteristic is referred to as the protogyny phenotype. Here, we aimed to elucidate the mechanism underlying the protogyny phenotype in the fruit fly using our newly developed *Drosophila* Individual Activity Monitoring and Detecting System (DIAMonDS), which can continuously detect the precise timing of both pupariation and eclosion of individual flies. Via this system, following the laying of eggs, we detected the precise time points of pupariation and eclosion of a large number of individual flies simultaneously and succeeded in identifying the tiny differences in pupal duration between females and males. We first explored the role of physiological sex by establishing transgender flies via knockdown of the sex-determination gene, *transformer* (*tra*) and its co-factor *tra2*, which retained the protogyny phenotype. In addition, disruption of dosage compensation by *male-specific lethal* (*msl-2*) knockdown did not affect the protogyny phenotype. The *Drosophila* master sex switch gene—*Sxl* promotes female differentiation via *tra* and turns off male dosage compensation through the repression of *msl-2.* However, we observed that stage-specific whole-body knockdown and mutation of *Sxl* induced disturbance of the protogyny phenotype. These results suggest that an additional, non-canonical function of *Sxl* involves establishing the protogyny phenotype in *D. melanogaster.*

**Author summary:** A wide variety of animals show differences in time points of sexual maturation between sexes. For example, in many mammals, including human beings, females mature faster than males. This maturation often takes several months or years, and precisely detecting the time point of maturation is challenging, because of the continuity of growth, especially in mammals. Moreover, the reason behind the difference in sexual maturation time points between sexes is not fully understood. The fruit fly *Drosophila*—a model organism—also shows biased maturation between the sexes, with females emerging 4 h faster than males (a characteristic known as the protogyny phenotype). To understand the mechanism underlying the protogyny phenotype, we used our newly developed system, *Drosophila* Individual Activity Monitoring and Detecting System (DIAMonDS), to detect the precise eclosion point in individual fruit flies. Surprisingly, our analysis of transgender flies obtained by knockdown and overexpression techniques indicated that a physiological gender might not be necessary requirement for protogyny and that a non-canonical novel function of the fruit fly master sex switch gene, *Sxl*, regulates protogyny in fruit flies.

## Introduction

The time taken to reach sexual maturity is often unequal between the sexes of numerous animal species; protogyny refers to the phenotype characterized by females maturing first and protandry refers to the phenotype characterized by males maturing earlier than females [1,2]. A search of the AnAge database (https://genomics.senescence.info/species/) revealed that approximately one-third of animals throughout the animal kingdom show sexual dimorphism in sexual maturation timing, among which poikilotherms tend to exhibit protandry, whereas homeotherms tend to exhibit protogyny (S1 Table) [3]. Although there is minimal information on arthropods in the AnAge database, several reports have indicated that male adults tend to emerge somewhat earlier than females for many insect species [2, 4–6]. Numerous hypotheses have been proposed to explain the strategy of protogyny and protandry with respect to increasing fitness; however, the detailed mechanism or benefit remains to be elucidated [1, 6–8].

To better understand the evolutionary significance of the sex bias in the sexual maturation time point, it is also important to elucidate the molecular mechanism underlying the protogyny and protandry phenotypes. However, these molecular aspects remain unclear, mainly owing to difficulties in precisely measuring the timing of maturation of individuals simultaneously and for a long period with available techniques.

In the fruit fly *Drosophila melanogaster*, adult females emerge quickly, before males (protogyny phenotype), with only a 4-h difference in eclosion timing [9]. Therefore, *D. melanogaster* offers a potentially useful model to elucidate the molecular mechanism underlying the sexual dimorphism in sexual maturation. We established a new system, *Drosophila* Individual Activity Monitoring and Detection System (DIAMonDS), which can automatically detect the phase-conversion timing of individual flies, such as the timing of pupariation, adult eclosion, and death, with high temporal resolution [10]. DIAMonDS enables time-lapse- and multi-scanning to simultaneously determine the time points of pupariation and eclosion in a large number of individuals under several chemical and environmental conditions and against different genetic backgrounds. Using DIAMonDS, we could precisely detect the 4-h difference of eclosion timing between sexes and further revealed that this was solely due to a difference in pupal duration [10].

In this study, we further applied DIAMonDS to evaluate the genetic regulation of the protogyny phenotype of *D. melanogaster.* As fruit flies alter their developmental rates when exposed to different environmental conditions [11–13], we first explored the effect of temperature and nutrients on the protogyny phenotype. We next manipulated the *transformer (tra)* gene and its co-factor *transformer-2 (tra2)—*which play essential roles in determining the physiological sex of cells—to change the sex of the flies and evaluated the effect on the protogyny phenotype. Sex chromosome dosage compensation is also regulated differentially by sex, and the male-specific lethal complex is a key player in the dosage compensation machinery in *Drosophila* [14, 15]. Therefore, we also investigated the possibility that the dosage compensation pathway contributes to the protogyny phenotype by knocking down expression of the *male-specific lethal* (*msl*) *2* gene. Finally, we evaluated the potential role of the Sex-lethal gene *(Sxl)*, which encodes an RNA splicing enzyme and acts as a master regulator of the sex-determination pathway [16–18] and also regulates the expression of its downstream genes—*tra* and *msl-2.*

## Results

### Environmental stability of the protogyny phenotype in *D. melanogaster*

The sexual difference in pupal duration was maintained under a high temperature condition of 29°C (Fig 1A) and was also not altered under several nutritional conditions, including various sugar and yeast concentrations (Fig 1B and C). These results indicated that the sexual difference of pupal duration is a very stable phenotype in *D. melanogaster.* Therefore, we further used this difference to evaluate the molecular and genetic aspects underlying protogyny.

**Fig 1.**
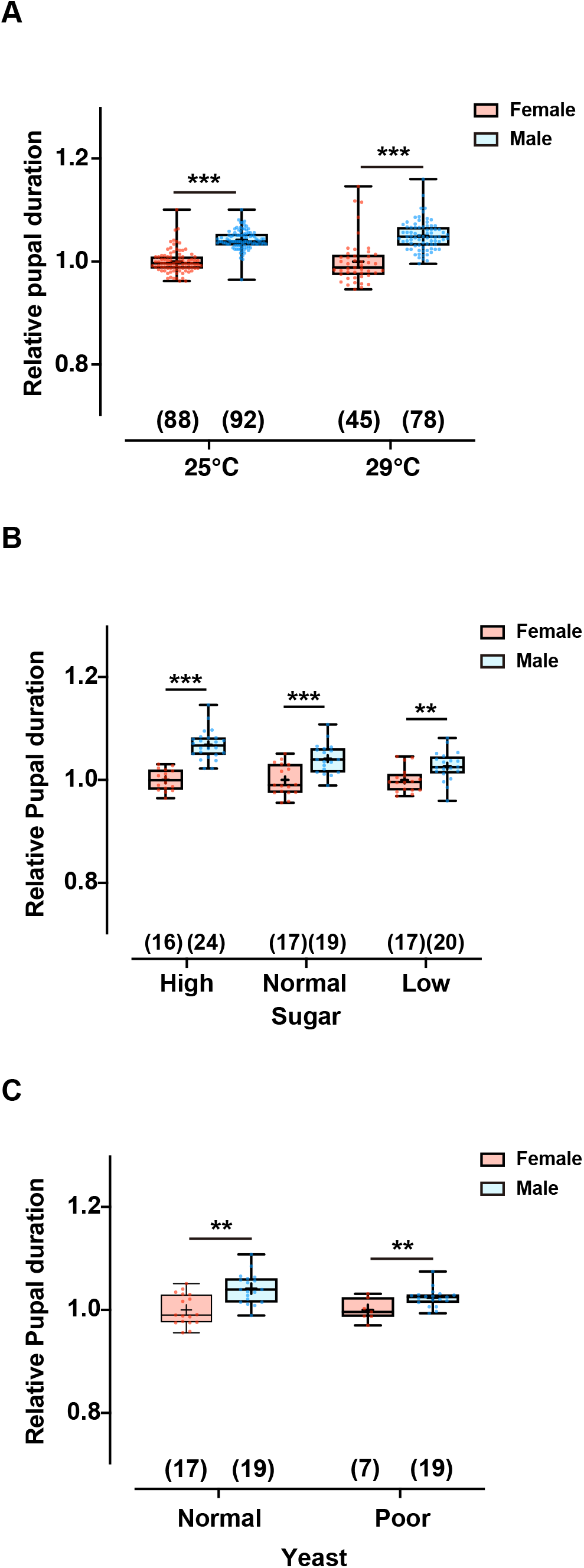
Environmental stability of the protogyny phenotype. A. Effect of rearing temperature of 25°C and 29°C. B. Effect of sugar concentration in the media. High, normal, and low sugar media contain 1 M, 0.15 M, and 0.05 M sugar, respectively, in addition to the other components of the normal medium. C. Effect of yeast concentration in the media. The poor yeast medium contains one-third the yeast concentration of normal fly medium. The number of flies analyzed is indicated in parentheses on each graph. Whiskers indicate minima and maxima (\*\*\**p* < 0.001; \*\**p* < 0.01; Student’s unpaired *t*-test).

### Forced sex change does not affect the protogyny phenotype based on the genotype

A previous study [19] revealed that *tra2* knockdown or *tra* overexpression in the whole body induced a sex transformation so that the phenotypic sex was opposite to the genotypic sex, which also altered body size (Fig 2A). We confirmed that the phenotypic sex transformation of *D. melanogaster* can be controlled by genetic manipulation of *tra* or *tra2* expression independent of the sexual genotype using *UAS-tra2* RNA interference (RNAi)-mediated knockdown or *UAS-traF* overexpression with ubiquitous *GAL4* drivers (Fig 2B). Pupal durations were then compared between siblings with XX and XY genotypes, respectively (S1 Fig). The phenotypic transformation induced by *tra2* knockdown or *traF* overexpression did not alter the sexual difference of pupal duration based on the chromosomal sex (Fig 2C-D). These results suggested that phenotypic sex is not critical for the protogyny phenotype, which is also independent of the *tra/tra2* pathway.

**Fig 2.**
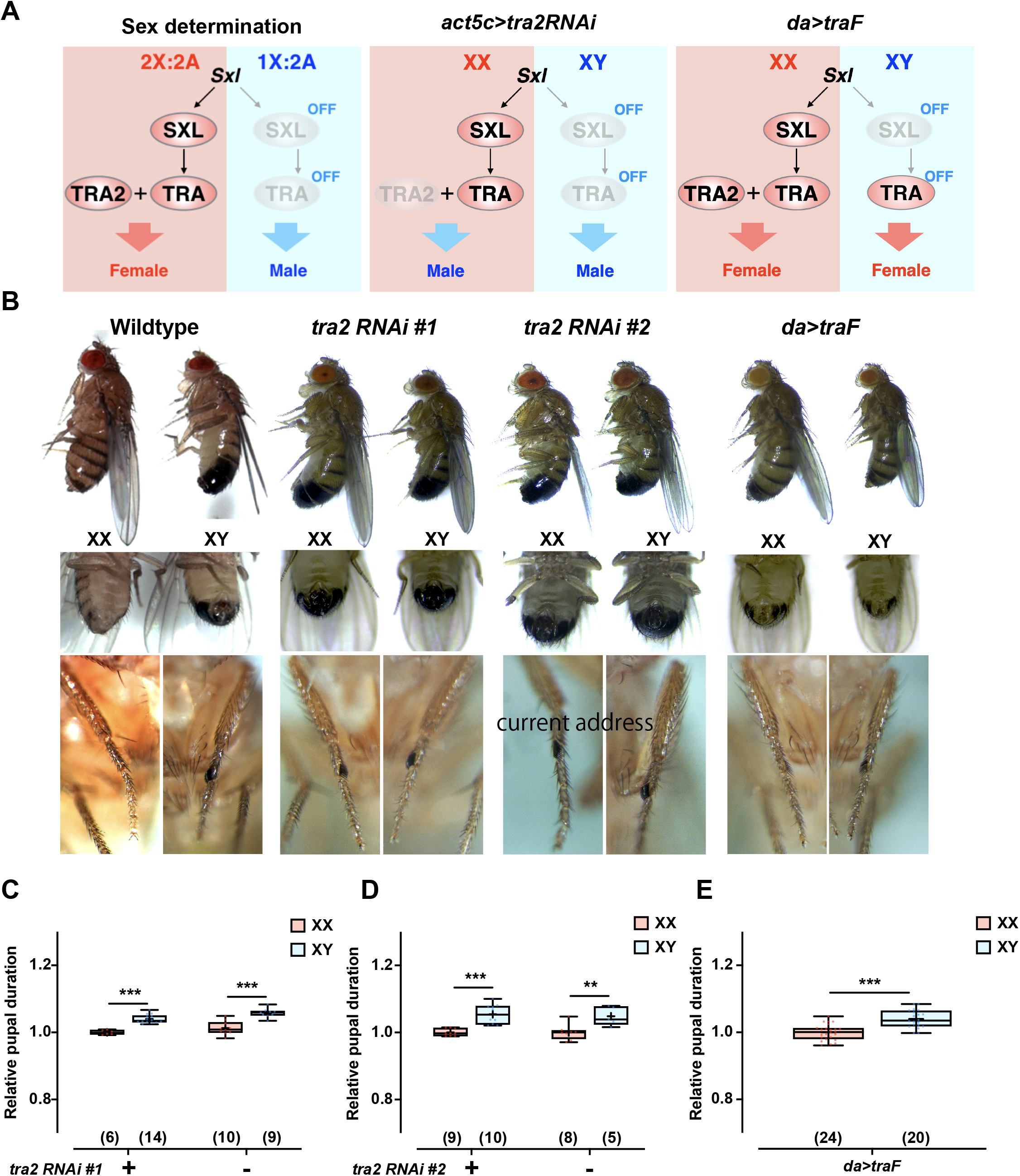
Alteration of *tra2* and *tra* expression does not affect the protogyny phenotype. A. Schematic presentation of the sex-determination pathway and effect of alteration of *tra2* or *traF* expression. B. Photographs of external morphological sexual traits of wild-type, *act5c>tra2* RNAi #1, *act5c*>RNAi #2, and *da>traF* adults. C–E. Effect of *act5c>tra2* RNAi #1 (C), *act5c*>RNAi #2 (D), and *da>traF* adults (E) on the protogyny phenotype. The number of flies analyzed is indicated in parentheses on each graph. Whiskers indicate minima and maxima (\*\*\**p* < 0.001; \*\**p* < 0.01; Student’s unpaired *t*-test).

### Disturbance of the dosage compensation pathway could not alter the protogyny phenotype

Dosage compensation machinery is not assembled in *Drosophila* females, because *msl-2*, a key gene of assembly of the MSL complex is not translated. [14, 15, 20]. Thus, next, we investigated the possibility of contribution of the dosage compensation pathway to development of the protogyny phenotype.

Ubiquitous knockdown of *msl-2* (Fig 3A) successfully induced male-specific semi-lethality (Fig 3B), which in turn reduced the *msl-2* expression level in males (Fig 3C). However, *msl-2* knockdown did not change the sexual difference of pupal duration, suggesting that the sex chromosome dosage compensation machinery does not commit to the protogyny phenotype (Fig 3D and E and S2 Fig).

**Fig 3.**
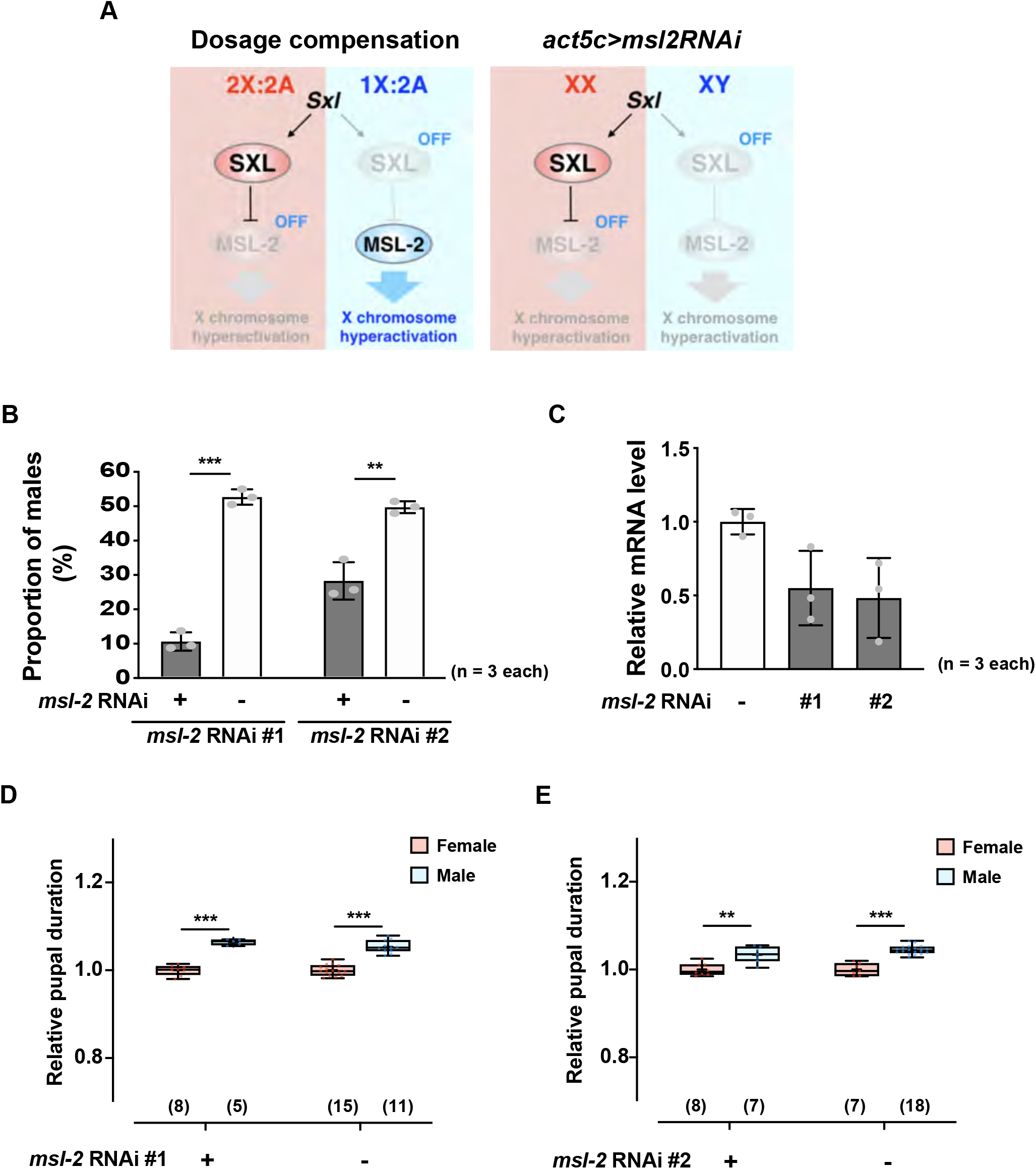
Alteration of *msl-2* expression does not affect the protogyny phenotype. A. Schematic presentation of the dosage compensation pathway and effect of *msl-2* expression alteration. B. Proportion of eclosed males of *act5c>msl-2 #1* and *act5c>msl-2 #2* lines. C. Relative *msl-2* mRNA level of the *act5c>msl-2 #1 and act5c>msl-2 #2* groups. D-E. Effect of *act5c>msl-2 RNAi #1* (D) and *act5c>msl-2 RNAi #2* (E) on the protogyny phenotype. The number of flies analyzed is indicated in parentheses on each graph. Whiskers indicate minima and maxima (\*\*\**p* < 0.001; \*\**p* < 0.01; Student’s unpaired *t*-test).

### The protogyny phenotype is determined in an *Sxl-*dependent manner

Ubiquitous *Sxl* knockdown using *act5c-GAL4* was not successful owing to its lethal phenotype. Therefore, we further attempted pan-neuronal *Sxl* knockdown using *elav-GAL4*, which did not influence the sexual difference in pupal duration (S3 Fig). To avoid lethality during larval development, we used a gene-switch system, which can induce GAL4 by administrating the glucocorticoid receptor antagonist RU486 [21]. F1 larvae were derived from parents of a *UAS-Sxl* RNAi transgenic fly and an *act5c-GS-GAL4* fly reared in normal condition, and early 3rd-instar larvae were transferred to a 96-well-microplate containing media with or without the RU486. Some adult F1 females that escaped from the RU486-containing media showed partial sexual transformation morphologically, indicating that RU486-dependent transformation succeeded in this condition (S4 Fig). Moreover, only the F1 flies reared in RU486-containing media did not exhibit the sexual difference of pupal duration, suggesting that *Sxl* might regulate the protogyny phenotype (Fig 4A).

**Fig 4.**
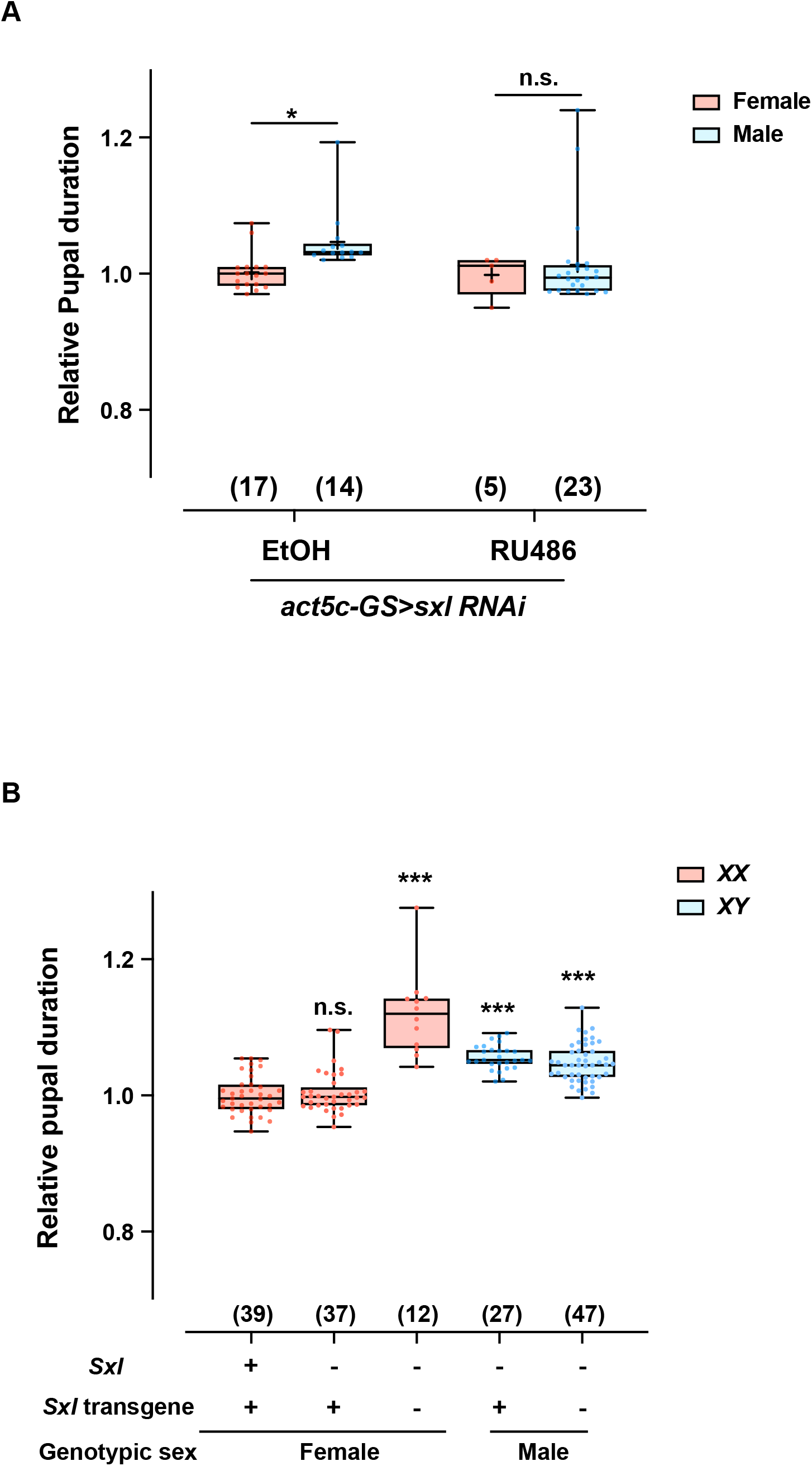
Alteration of *Sxl* expression affects the protogyny phenotype. A. Effect of *act5c-GS>Sxl* RNAi on the protogyny phenotype of flies grown in media with or without RU486. B. Effect of *Sxl* mutation on the protogyny phenotype in flies with and without the *Sxl* transgene. The number of flies analyzed is indicated in parentheses on each graph. Whiskers indicate minima and maxima (\*\*\**p* < 0.001; \**p* < 0.05; n.s., no significant difference; Student’s unpaired *t*-test).

To confirm this interpretation, we also tried to use trans-heterozygous *Sxl^M1,Δ33^/Sxl^f7,M1^* masculinized females [22–24], which have low viability but exhibit the same revertant eclosion to the adult stage [22]. Using our cross scheme, we were able to produce two genotypes of *Sxl^M1,Δ33^/Sxl^f7,M1^* (*Sxl^-^*) flies with and without an extra *Sxl* transgene (Fig 4B). The *Sxl^-^* flies without an extra *Sxl* transgene showed a significantly longer pupal duration in comparison with that of *Sxl^+^* females, reaching the same length as that of the male flies. The phenotype of *Sxl^-^* flies completely recovered by introduction of the extra *Sxl* transgene (Fig 4B).

## Discussion

In this study, we applied our recently developed DIAMonDS to explore the molecular mechanism underlying the very small but consistent sex difference in eclosion timing due to a difference of pupal duration.

Many morphological and physiological traits exhibit a sex difference, which may be controlled by a canonical sex-determination pathway [25]. However, the protogyny phenotype was not disturbed in genetically induced transgender flies established by controlling *tra* or *tra2* gene expression or by knockdown of *msl-2.* These results suggest that a morphological or physiological (dosage compensation) sex difference does not play a central role in controlling the protogyny phenotype, as manipulating these factors did not influence the length of male pupal duration. However, further genetic manipulation experiments demonstrated that the non-canonical function of *Sxl* regulates the eclosion timing and produces the protogyny phenotype in *D. melanogaster*, as females with loss-of-function mutations or knockdown of *Sxl* exhibited a pupal period of the same length as that of males.

*Sxl* expression is activated in the presence of two X chromosomes in female early embryos, and is maintained via positive auto-regulation [16, 22, 26]. Sxl also regulates splicing of its downstream components, including *tra* and *msl-2*, which play crucial roles in the sex-determination cascade and dosage compensation, respectively [27, 28]. Therefore, our results suggest that recently identified non-canonical Sxl pathways could be involved in the protogyny phenotype.

Indeed, Sxl protein has been suggested to interact with other targets. *Nanos (nos)* RNA can bind directly with Sxl protein in ovarian extracts, and loss-of-function studies suggested that Sxl enables the switch from germline stem cells to committed daughter cells through *nos* post-transcriptional down-regulation [29]. Sxl protein can also bind with *Notch* (*N*) mRNA and appears to negatively control the *N* pathway [30]. Genome-wide computational screening for Sxl targets also identified an ATP-dependent RNA helicase, Rm62, as a novel potential target [31]. Rm62 was inferred to be involved in alternative splicing regulation and is required for the RNAi machinery [32, 33]. A pan-neuronal RNA-binding protein of the ELAV family, *found in neurons (fne)*, was also shown to be downregulated by *Sxl* in female heads, independent of *tra/tra2* regulation [34]. Sxl can enhance nuclear entry of the full-length Cunitus interuptus (Ci) protein, suggesting a contribution to the sex difference in growth rate, although their physical interaction has not been confirmed [35]. However, there is no evidence that these non-canonical targets of Sxl directly affect eclosion timing. Therefore, further studies are required to demonstrate whether these Sxl targets, or another novel target, could contribute to the protogyny phenotype.

In this study, conditional knockdown of *Sxl* in the nervous system, fat body, or intestinal stem cells and enteroblasts did not disrupt the protogyny phenotype; only *Sxl* knockdown induced in the whole body could induce the delayed eclosion in females. As knockdown of *Sxl* from the early developmental stage strongly affected viability, this toxicity might be one reason for our inability to identify the responsible organ or tissue that regulates the protogyny phenotype in this study.

The independence of the protogyny phenotype from the canonical sex-determination pathway is very intriguing with respect to understanding evolution of the sex difference in sexual maturation. *Sxl* does not appear to play a role in sex determination in most insects [24, 36–38]. Several reports indicated that orthologs of *Sxl* had no sex-determinant role in *non-Drosophila* species, including in *Diptera* [37, 39, 40]. In Drosophilidae, ancestral *Sxl* was duplicated to *Sxl* and *sister of sex lethal* (*ssx*); the new *ssx* gene plays a role of ancestral *Sxl*, suggesting that *Sxl* might have evolved to function as a novel sex-determinant gene in Drosophilidae [38]. A detailed phylogenetic study revealed that a male-specific exon, and likely embryo-specific exon, originated after the divergence between the Drosophilidae and Tephritidae families but before the split of the *Drosophila* and *Scaptodrosophila* genera [41]. We presume that the implementation of *Sxl* in the sex-determination pathway may be significantly involved in acquisition of the protogyny phenotype in *Drosophila.* Therefore, we expect that identification of the target of non-canonical *Sxl* sex-specific regulation for the protogyny phenotype might help to promote understanding of the evolutionary aspects of protogyny.

## Material and methods

### *Drosophila* stocks

All flies were maintained at 25°C on standard laboratory medium as described previously [42]. The following stocks were obtained from the Bloomington *Drosophila* stock center (BDSC): *w^1118^* (wild-type; BDSC 5905), *act5c-GAL4* (BDSC 3954)*, da-GAL4* (BDSC8641)*, elav-GAL4* (BDSC 458)*, elav-GAL4; UAS-dcr-2* (BDSC 25750)*, P{CaryP} attP2* (BDSC 36303)*, UAS-tra2 RNAi #1* (BDSC 56912)*, UAS-tra2 RNAi #2* (BDSC 28018), *UAS-traF* (BDSC 4590)*, UAS-msl-2 RNAi #1* (BDSC 31627)*, UAS-msl-2 RNAi #2* (BDSC 35390)*, UAS-Sxl RNAi #1* (BDSC 34393)*, UAS-Sxl RNAi #2* (BDSC 38195), *Sxl^f7,M1^; P{Sxl.+tCa}9A/+* (BDSC 58486), and *Sxl^M1,fΔ33^/Binsinscy* (BDSC 58487). Three gene-switch *Gal4* driver lines, *act5c-GS-GAL4, S106-GS-GAL4*, and *5961-GS-GAL4*, were a kind gift from Dr. Akagi [43].

### Measurement of pupal duration

We used our recently developed DIAMonDS to measure pupa duration at the individual level. The wandering 3^rd^-instar larvae were collected from rearing vials, and a single larva was placed in the well of a 96-well microplate with normal medium. The plate was then placed on a flatbed CCD scanner to obtain time-lapse images until all flies were eclosed. The time-lapse image dataset was then analyzed using Sapphier software as described previously [10].

To compare the effect of *Sxl* mutation on pupal duration, *Sxl^f7,M1^; P{Sxl.+tCa}9A/+* females were crossed with *w^1118^/Y* males. The F1 progeny *Sxl^f7,M1^/Y; P{Sxl.+tCa}9A/+* males were then crossed with *Sxl^M1,fΔ33^/Binsinscy* females. Each genotype of the F2 flies was then assessed for pupal duration using the DIAMonDS. To induce the gene-switch *Gal4* driver, RU486 (Mifepristone, Sigma, St. Louis, MO, USA) reagent was dissolved in ethanol and added to the medium at a final concentration of 100 μg/ml.

To detect the sex genotype of the flies, genomic DNA was extracted from single adults by homogenization in 50 μl of squishing buffer (10 mM Tris-HCl [pH 8.2], 1 mM EDTA, 25 mM NaCl, 200 μg/ml proteinase K), and incubated at room temperature for 20 min, followed by inactivation at 95°C for 5 min. The extracted genomic DNA was subjected to polymerase chain reaction (PCR) analysis using a WD repeat-containing protein on Y chromosome *(WDY)-* and *Rp49-specific* primer mix by ampliTaq Gold 360 master mix (Applied Biosystems, Foster City, CA, USA), and then the amplified DNA fragments were separated by 2% agarose gel electrophoresis (S1 Fig).

### Reverse transcription-quantitative PCR (RT-qPCR)

Total RNA was extracted from the whole adult body (for measuring *msl-2* expression knocked down by *act5c-GAL4)* and from the dissected larval central nervous system (for measuring *Sxl* expression knocked down by *elav-GAL4)* using Isogen II (Nippon Gene, Tokyo, Japan), and then RT-qPCR was performed using a One Step SYBR PrimeScript PLUS RT-PCR kit (Takara Bio, Shiga, Japan) and Applied Biosystems ABI Prism 7000 Sequence Detection System. All mRNA expression levels were normalized to the levels of *rp49* mRNA. We used the following primers for qPCR (5’-3’) for detecting *msl-2* and *Sxl* mRNA levels: *Sxl*, forward primer (5’-CCAATCTGCCGCGTACCATA-3’), reverse primer (5’-AATGGAACCGTACTTGCCGA-3’); *msl-2*, forward primer (5’-CACTGCGGTCACACTGGCTTCGCTCAG-3’), reverse primer (5’-CTCCTGGGCTAGTTACCTGCAATTCCTC-3’); and *rp49*, forward primer (5’-GATGACCATCCGCCCAGCATAC-3’), reverse primer (5’-AGTAAACGCGGTTCTGCATGAGC-3’).

### Statistical analysis

All data were analyzed and graphs were plotted using Prism version 8 (GraphPad Software, San Diego, CA, USA). Data are presented as means ± standard deviation. Student’s unpaired two-tailed *t*-test was performed to compare differences between two groups in each experiment, and Dunnett’s one-way analysis of variance was used for multiple comparisons; p < 0.05 was considered to indicate a statistically significant difference.

## Acknowledgments

The authors are deeply grateful to Tadashi Uemura (Kyoto University) for the helpful support. They also thank Yuko Iijima and Taishi Matsumura for their technical assistance. We received generous support from Yoichi Shinkai (RIKEN CPR) and Shunsuke Ishii (RIKEN CPR). This work was also supported by the AMED (Japan Agency for Medical Research and Development) [JP18gm1110001 to KHS. and SK].

## Supporting information

**S1 Fig. Detection of the sex genotype of single flies.** A. WDY primer sets and results of single-fly PCR for females and males of *w^1118^, act5c>tra2 RNAi #1, act5c>RNAi #2*, and *da>traF* fly lines. B. Results of single-fly PCR for *act5c>tra2 RNAi #1.* C. Results of single-fly PCR for *da>traF.*

**S2 Fig. Effect of *da>msl-2 RNAi #1* and *da>msl-2 RNAi #2* on the protogyny phenotype.** The number of flies analyzed is indicated in parentheses in each graph. Whiskers indicate minima and maxima (\*\*\**p* < 0.001; Student’s unpaired *t*-test).

**S3 Fig. Alteration of *Sxl* expression in the central nervous system does not affect the protogyny phenotype.** A, B. Relative *Sxl* mRNA levels of the *elav>Sxl RNAi #1/#2; dcr2* (A) and *elav>Sxl RNAi #1/#* (B) fly lines. C. Effect of *elav>Sxl RNAi #1* and *elav>Sxl RNAi #2* on the protogyny phenotype. D. Effect of *S106-GS>Sxl RNAi #1* and *5961-GS>Sxl RNAi #2* on the protogyny phenotype. The number of flies analyzed is indicated in parentheses on each graph. Whiskers indicate minima and maxima (\*\*\**p* < 0.001; \*\**p* < 0.01; \**p* < 0.05; n.s., no significant difference; Student’s unpaired *t*-test).

**S4 Fig. Photographs of external morphological sexual traits of *act5c-GS>Sxl RNAi #1* flies in medium with and without RU486.**

**S1 Table.** Abundance of Protandry and Protogyny in vertebrate species

| **Class** | **Protandry** | **Protogyny** | **No bias** | **Total (number of species)** |
| --- | --- | --- | --- | --- |
| **Poikilotherm** |  |  |  |  |
| Chondrichthyes | 43 | 8 | 10 | 61 |
| Teleostei | 58 | 27 | 150 | 235 |
| Amphibia | 20 | 2 | 56 | 78 |
| Reptilia | 14 | 1 | 55 | 70 |
| **Homeotherm** |  |  |  |  |
| Aves | 12 | 21 | 645 | 678 |
| Mammalia | 77 | 233 | 264 | 574 |
| **Total** | 224 | 292 | 1180 | 1696 |

Calculated from dataset of AnAge Database (https://genomics.senescence.info/species/)

**Figure 2-Supplementary figure 1.**
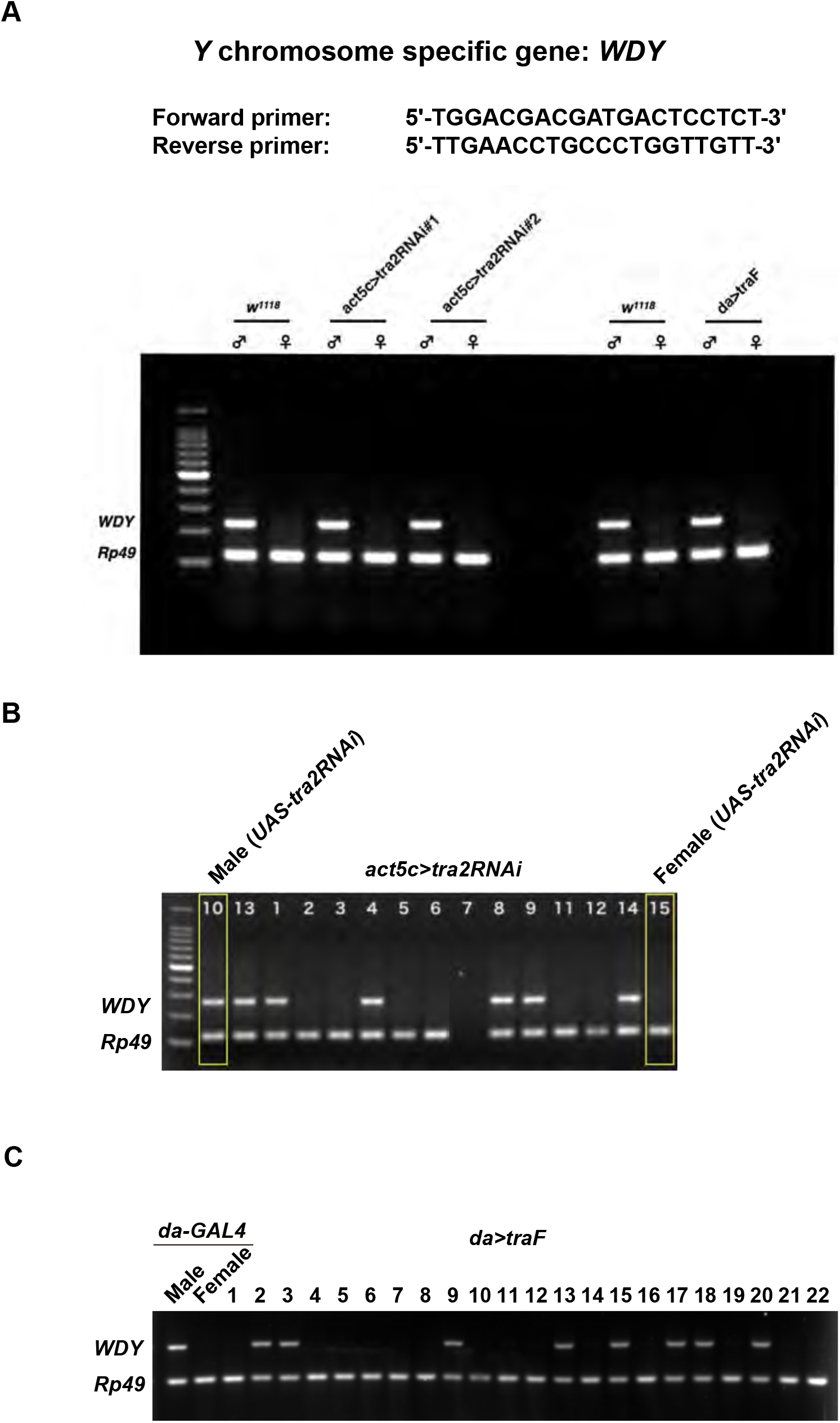

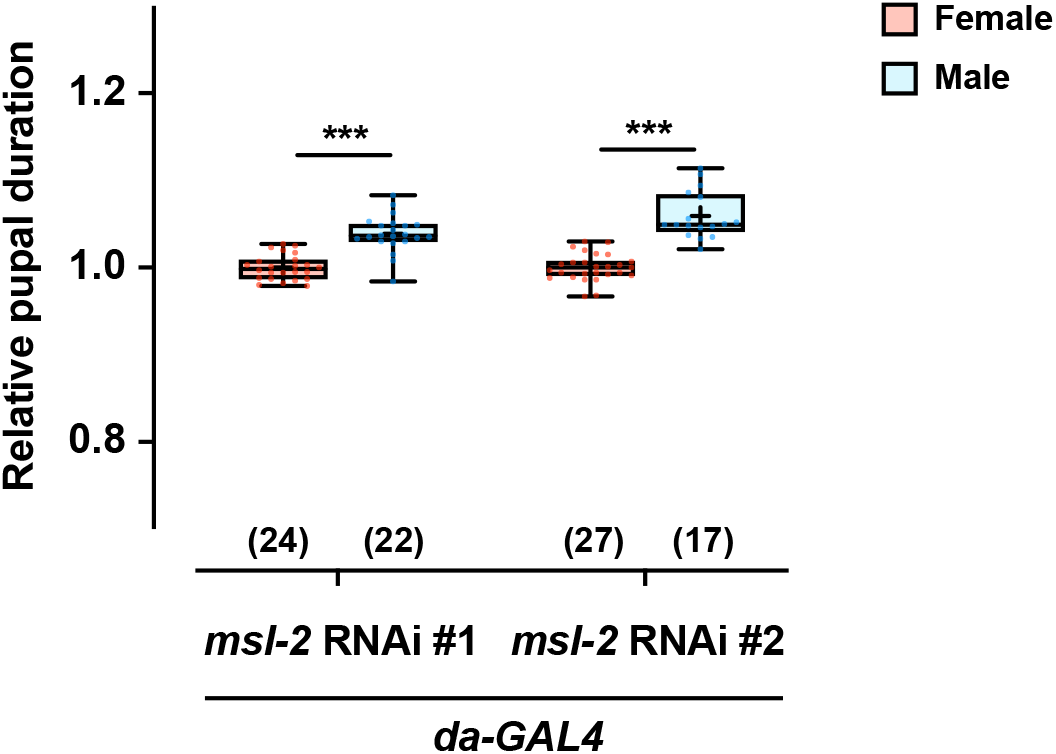

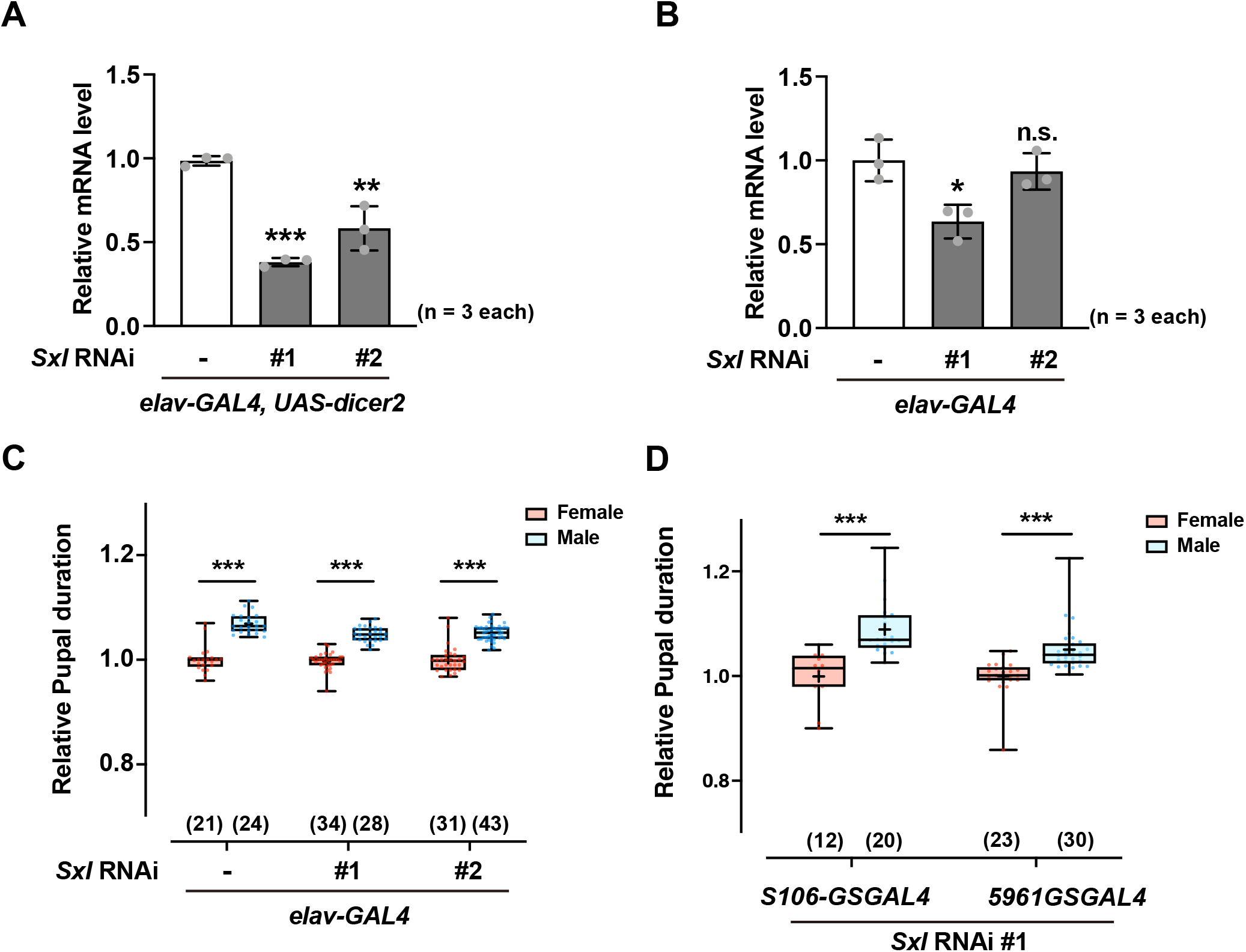

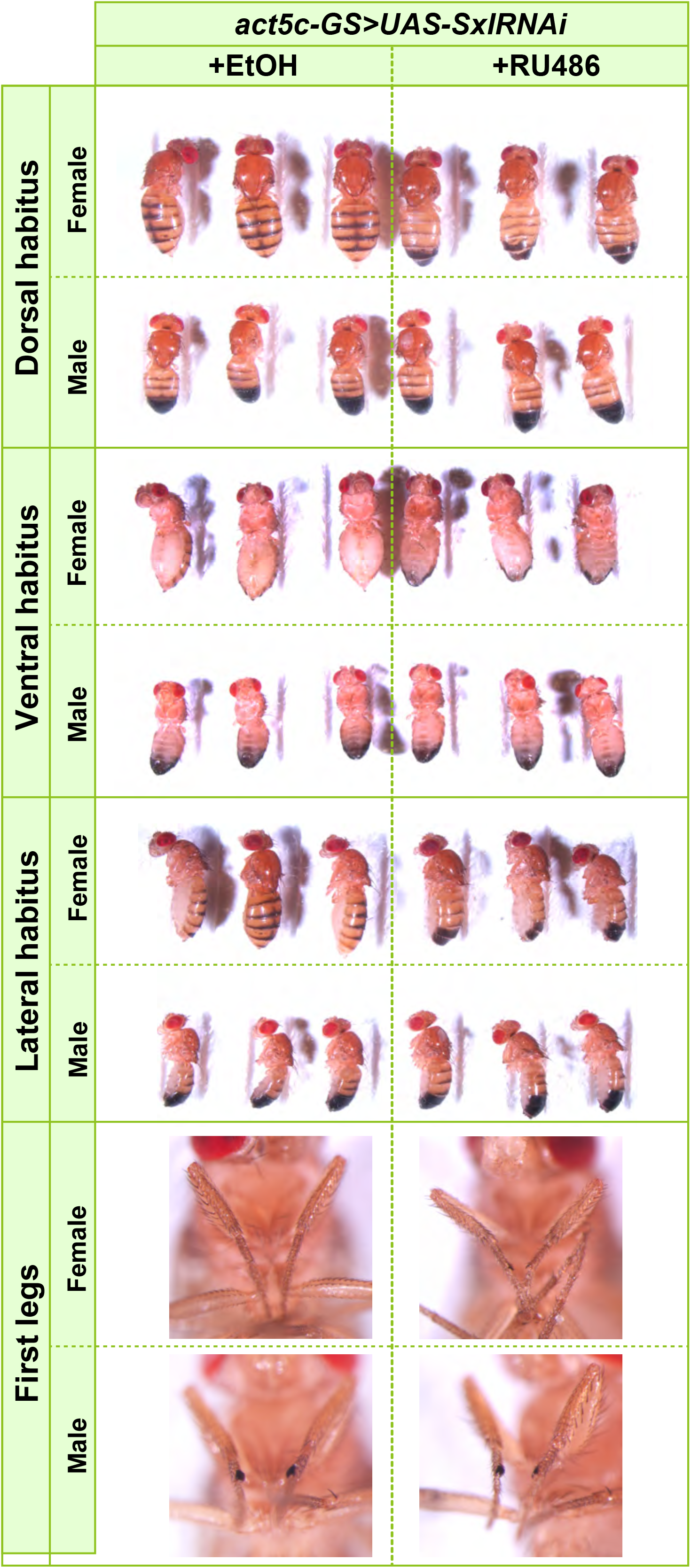

